# Frontostriatal salience network expands as executive networks contract in Obsessive-Compulsive Disorder

**DOI:** 10.1101/2025.05.21.654808

**Authors:** Matilde M. Vaghi, Martin Norgaard, Sunjae Shim, Jaime Ali H. Rios, Patrick G. Bissett, Carolyn I. Rodriguez, Russell A. Poldrack

## Abstract

Obsessive-compulsive disorder (OCD), marked by intrusive thoughts (obsessions) and repetitive behaviours (compulsions), is linked to dysfunction in frontostriatal circuits. However, neural differences potentially contributing to these alterations are often small, and conflicting evidence obscures the directionality and underlying mechanisms of these alterations. Like many psychiatric conditions, OCD follows a fluctuating symptom trajectory, with symptoms shifting dramatically over months—either naturally or due to treatment. Yet, the absence of longitudinal neuroimaging studies limits our understanding of the neural mechanisms driving these changes.

Here, we used precision functional mapping in highly sampled individuals with OCD, uncovering a striking imbalance within frontostriatal networks. We identified a twofold expansion of the salience network in these individuals and a concomitant contraction of the frontoparietal network. Notably, salience network expansion was driven by border shifts, encroaching on adjacent executive networks and leading to their contraction. This imbalance within frontostriatal networks may relate to excessive attention to internal stimuli and lack of goal-directed control, which are classically observed in OCD. Longitudinal analyses of neuroimaging data collected over several months revealed frontostriatal connectivity changes tracking symptom severity. Overall, these findings pinpoint network-level features that may confer risk for individuals with OCD and highlight dynamic connectivity shifts tied to severity of OCD symptoms over time. By isolating frontostriatal abnormalities at the individual level in OCD and their relationship to symptom severity for the first time, this work paves the way for more targeted, personalized treatment strategies and identifies precision functional mapping as a model for precision psychiatry.

## Main

Obsessive-compulsive disorder (OCD) is a major cause of disability worldwide and one of the most well-characterized neurocircuit-based psychiatric disorders^1–6^. It is hypothesized to arise from dysregulation in frontostriatal circuits, which integrate motor, limbic, and cognitive pathways to support emotional regulation, goal-directed behaviour, and habit formation during incentive-based learning^7–9^. However, despite the broad appeal of this hypothesis, the precise nature of frontostriatal imbalances in OCD has remained elusive. The neurobiological mechanisms underlying hallmark OCD symptoms— such as intrusive thoughts with a strong emotional load and compulsive behaviours that often become habitual and resistant to goal-directed control—are poorly understood. Furthermore, little is known about the neural mechanisms that drive symptom fluctuations over time, whether naturally occurring or in response to treatment.

A key problem with current research is that it has focused only on case-control comparisons at a single time point, neglecting the importance of individual level analysis. The traditional cross-sectional focus also provides limited insight into whether observed effects are stable over time or state-dependent. More generally, reliance on group-level and cross-sectional inference based on a relatively small amount of data per participant has hindered biomarker research using functional magnetic resonance imaging (fMRI) yielding inconsistent results and suggesting that the typical study design may not be ideal for identifying actionable targets for personalized diagnosis and intervention.

Recent advances offer a pathway toward individualized models of brain function. Study designs have been formalized entailing the study and analysis of a single participant or a series of individuals over repeated sessions. These smaller and focused studies can maximize effect sizes and allow within-subject comparisons^10–13^. The emphasis on intensive longitudinal assessment is aimed at collecting large amounts of data within an individual to maximize measurement reliability. This approach is also well suited to gain within-person mechanistic insights by establishing links between neuroimaging metrics and symptom course. Accordingly, by leveraging dense sampling of individuals, the emerging field of precision functional mapping (PFM) enables the characterization of functional networks at the individual level, moving beyond the one-size-fits-all approach of conventional fMRI studies. Pioneering work using PFM in healthy individuals has shown individual specific features in topology of functional areas and networks that deviates markedly from group-average descriptions^11,12,14,15^. Individual differences in network topology have been shown to be reliable^14,16^, stable^10–12,17^, and heritable^18,19^. Importantly, individual network topologies extend beyond cortical regions to striatal and subcortical areas^20,21^. Additionally, several lines of research suggest that idiosyncratic differences in brain network topography are behaviourally meaningful, being associated with individual differences in cognition and behaviour^12,14,22,23^. Therefore, the emerging literature highlights the value of PFM in psychiatric conditions that show high inter-individual symptoms variance and underscore its value in OCD, which is heavily dependent on alterations within frontostriatal circuits. A recent study has identified characteristic network topologies in individuals with depression and shown connectivity changes that tracked fluctuations of specific symptoms^24^. However, apart from this one recent study involving individuals with depression, these tools have not yet been widely applied in psychiatric clinical populations, and it is unknown what the functional network topologies in individuals with OCD are and if and to what extent they might be shared with those identified in depression.

Like other psychiatric conditions, symptoms in OCD can fluctuate significantly within an individual over time—not only in response to intervention, but even when they are not actively receiving treatment, as seen through repeated assessment across periods of time in longitudinal studies^25–27^. However, the cross-sectional approach adopted by conventional neuroimaging studies, involving data acquired at a single time point or before and after intervention, does not enable understanding of the mechanisms that mediate variation in symptoms either naturally or due to treatment. Long-term and high sampling studies involving intracranial electrophysiology after Deep Brain Stimulation (DBS) have started to reveal the mechanisms that govern symptom transition in individual patients with OCD^28,29^. However, in conventional fMRI studies data collection over extended periods of time has not been widely implemented. Accordingly, single-subject estimates remain constrained by technical limitations related to sensitivity to imaging artefacts and noisy estimates due to limited amount of data. Recent advancements, including multi-echo fMRI^30^ and large-scale per-subject data acquisition^10,12^, now enable the generation of highly reliable functional connectivity measures and individualized network maps.

Here we used state-of-the-art PFM tools in conjunction with a dense sampling longitudinal design and multi-echo fMRI to map the topology of functional networks in individuals with OCD. We uncovered a striking imbalance within frontostriatal networks in OCD patients, finding that the frontostriatal salience network is expanded by more than twofold in individuals with OCD. Together with frontostriatal salience network expansion, we also identified a contraction of the frontoparietal network involved in executive control. Further analyses revealed that the salience network in OCD encroached on and displaced neighbouring functional systems generally implicated in executive functions, which are known to be compromised in OCD patients^31^. Longitudinal analysis of highly sampled individuals showed that changes in frontostriatal connectivity between nodes of the salience network and the dorsal striatum tracked fluctuations of OCD severity. Our findings provide the first individual-specific functional network characterization of OCD and highlight the potential of PFM for uncovering disorder-specific and transdiagnostic mechanisms. By moving beyond group-averaged descriptions, this approach may open new avenues for precision psychiatry, facilitating the development of individualized biomarkers and targeted interventions.

### Highly sampled longitudinal neuroimaging in OCD

We used highly sampled longitudinal neuroimaging in seven individuals with OCD, each undergoing 180 min of multi-echo resting-state fMRI scanning across 10 distinct sessions (**Fig. 1a**). To capture symptom severity the Yale-Brown Obsessive Compulsive Scale (Y-BOCS)^32^, a gold-standard measure for OCD symptoms, was administered at every session. Across study visits, the average Y-BOCS score was 20.74 ± 4.53 (range: 10–29), reflecting symptom severity ranging from mild to severe. Detailed imaging, demographic, and clinical information for this sample is available in the Methods section.

**Fig. 1.**
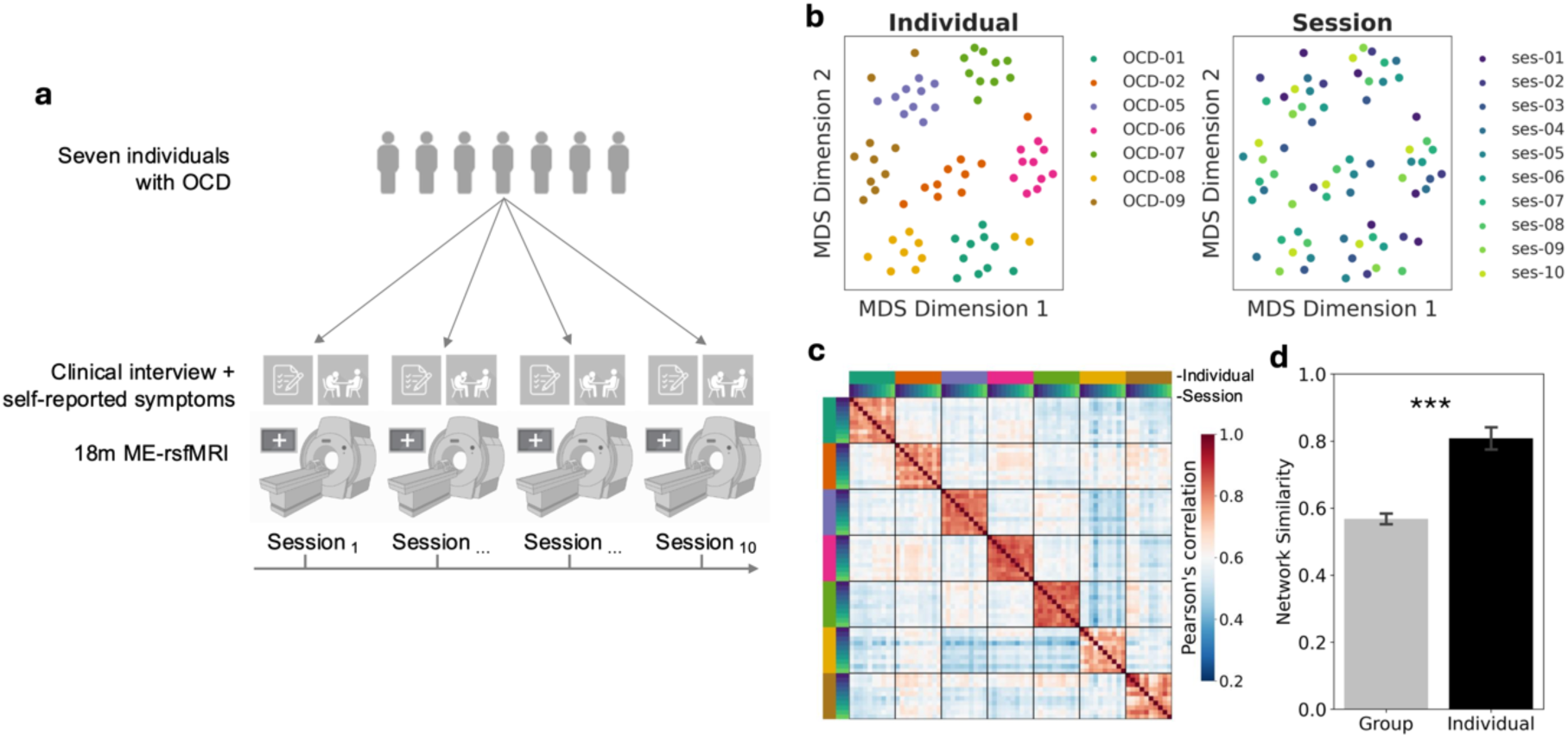
Individual identity dominates functional networks in highly sampled individuals with OCD. **a,** The experimental design involved repeated multi-echo resting-state fMRI scans (ME-rsfMRI) and clinical assessments of seven individuals with OCD. For each individual, data were acquired on 10 separate sessions over long periods of time (i.e., 3 months). **b,** A data-driven multi-dimensional scaling was used to depict the effect of individual identity on similarity among functional networks. In these plots each point represents a single functional network, plotted in a multidimensional space based on the similarity among networks. Networks are colored based on participant identity of each functional network (left) or session (right). The first two dimensions explaining 37% of the variation in functional networks were dominated by subject identity, suggesting it contributes most to functional network variance. **c,** Quantification of the similarity among functional brain networks was obtained by building a similarity matrix where each cell represents the correlation between a pair of functional networks. The matrix is organized first by individuals and then by session showing a stronger diagonal (similarity among networks from the same individual) and weaker off-diagonal (similarity across subjects). **d,** Barplot shows increased similarity for networks from the same individual but different sessions Individual, diagonal in panel c) than for networks from different individuals and sessions (group, off-diagonal in panel c) (*T* = 18.84, *P* < 0.001, paired two-tailed *t*-test), highlighting the strong influence of individual identity on network similarity.

### Individual identity dominates functional networks

We employed a combination of approaches to disentangle the contributions of group-level and individual-specific effects to network similarity. First, a data-driven analysis demonstrated that participant identity strongly influences network similarity (**Fig. 1b**). Functional networks from each individual and session were plotted in a multidimensional space based on their similarity (Euclidean distance). The first two dimensions, which account for 37% of the variation in functional networks, were predominantly shaped by subject identity. Then, we quantified network similarity within and across individuals by correlating all pairs of functional networks. Networks from the same individual (squares on the diagonal in **Fig. 1c**) were significantly more similar than those from different individuals (off-diagonal in **Fig. 1c**) (*T* = 18.84, *P* < 0.001, **Fig. 1d**). While networks across individuals and sessions exhibited substantial similarity (Group effect: mean *z*(*r*) = 0.57), reflecting a shared common structure, networks within the same individual showed even greater similarity (Individual effect: mean *z*(*r*) = 0.81), highlighting the strong influence of individual identity on network similarity. These findings replicate previous results in healthy volunteers^17^ and motivate the use of state-of-the-art PFM tools to delineate topology of functional brain networks at the subject-level in individuals with OCD.

### Expansion of frontostriatal salience network in individuals with OCD

We used PFM to chart the topology of functional brain networks for each of our seven highly sampled individuals with OCD. To compare network topographies between individuals with OCD and healthy controls (HC), the same PFM analytical procedures were applied to data from highly sampled healthy participants. Further methodological details are provided in the Methods section.

Using a validated analysis pipeline^24^ we identified marked differences between individuals with OCD and HC with respect to the salience network, which was strikingly larger in individuals with OCD (**Figs. 2a, 2b**). In all seven OCD individuals the salience network showed a greater expansion compared to the average observed in HC. On average, the salience network occupied 70% more of the cortical surface in OCD individuals relative to the mean in HC (4.54% ± 0.97% of cortex in OCD vs 2.67% ± 1.69% of cortex in HC) (**Fig. 2c**, top). This difference gave rise to a large group-level effect (Cohen’s *d* = 1.27). We found an expanded salience network in OCD not only in the cortex but also in the striatum where the representation of the salience network was increased in OCD individuals (35.88% ± 10.88% of striatum in OCD vs 17.36% ± 8.75% of striatum in HC) (**Fig. 2c**, bottom) with a large group-level effect (Cohen’s *d* = 1.93). The findings of an expanded salience network not only in the cortex but also in the striatum are in line with the notion of a functional and anatomical relation between frontostriatal areas through interconnected loops in which the cortex projects to the striatum and the striatum projects back to cortex indirectly via the thalamus^7–9^. Both effects (cortical and striatal expansion of the salience network in OCD individuals) were robust to methodological variations as they replicated without the use of global signal regression (**Extended Data Fig. 1**). The effects remained statistically significant when controlling for possible confounding factors including sex, head motion, and age (**Extended Data Figs. 2 a-c**) and with or without correction for potential site- or scanner-induced bias (**Extended Data Figs. 3a, 3b**). The effects of salience network expansion, both cortical and striatal, were not driven by individuals with comorbid diagnosis, as their exclusion did not affect the significance of the findings (**Extended Data Figs. 4a, 4b**). Overall, this set of analyses show consistent and reproducible expansion of the frontostriatal salience network in individuals with OCD.

**Fig. 2.**
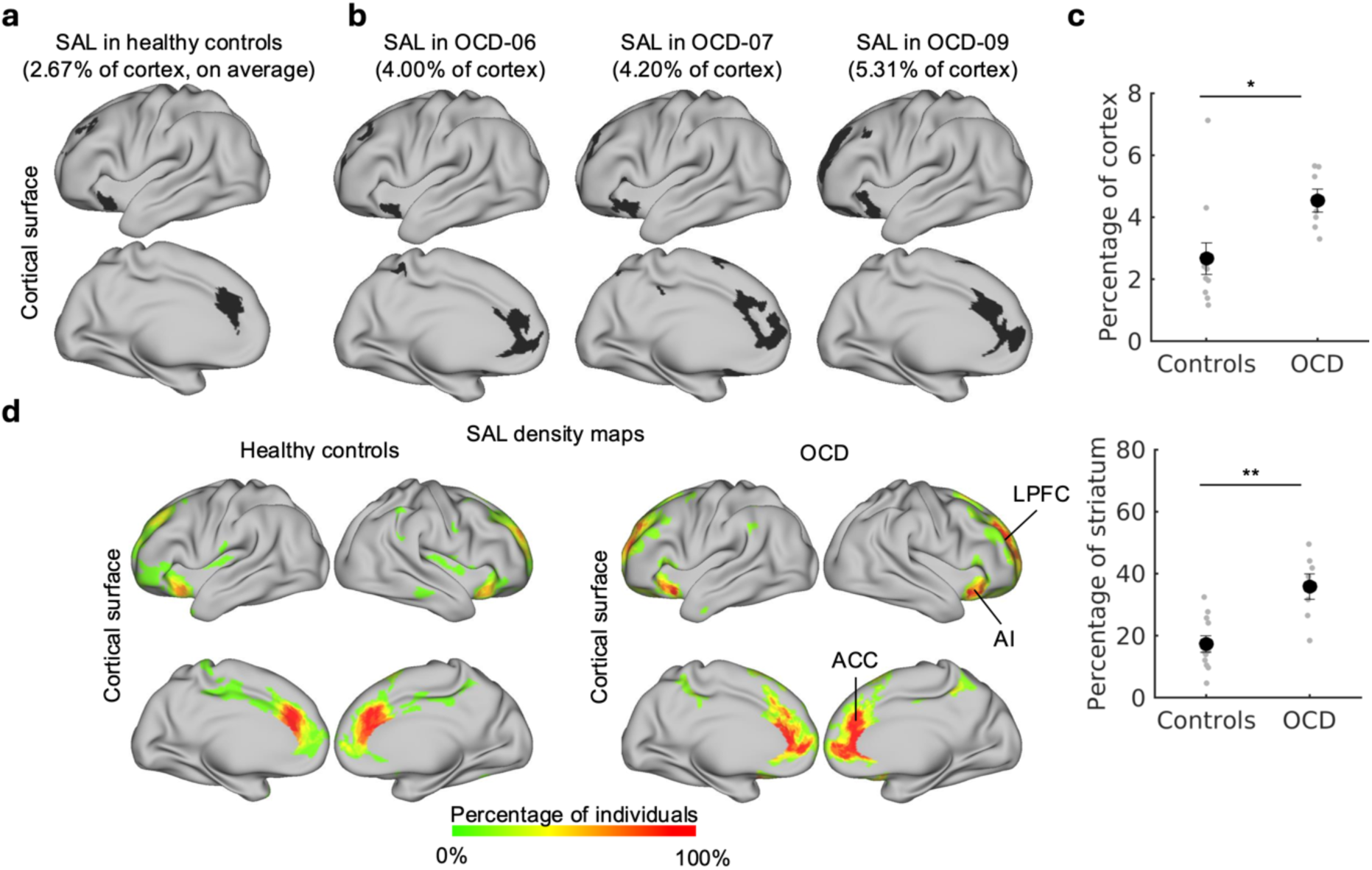
Cortical and striatal expansion of salience network in highly sampled individuals with OCD. **a,** The salience network (shown in black) covers an average of 2.67% of the cortical surface in HC, with nodes located in LPFC, ACC, and AI. **b**, The salience network in three representative individuals with OCD is displayed. In all OCD subjects, the salience network exhibits a greater expansion compared to the average observed in HC. **c,** On average, the salience network was 70% larger in individuals with OCD, and significantly expanded compared to HC as determined by a non-parametric Wilcoxon rank-sum *t*-test (*P* = 0.011, rank sum = 94, Cohen’s *d* = 1.27). Notably, the expansion of the salience network in OCD patients was also observed in the striatum (non-parametric Wilcoxon rank-sum *t*-test, *P* = 0.003, rank sum = 98, Cohen’s *d* = 1.93). Data are presented as mean ± s.d. **d,** Density maps confirmed that the spatial locations of salience network nodes were similar between HC and individuals with OCD. However, in OCD, the boundaries of the salience network extended further outward. ACC, anterior cingulate cortex; AI, anterior insular cortex; LPFC, lateral prefrontal cortex; OCD, obsessive-compulsive disorder; SAL, salience network.

Next, we used density maps to visualize the spatial organization of salience network nodes in individuals with OCD and HC. These quantified the percentage of individuals with salience network representation at each cortical vertex and they confirmed that locations of salience networks nodes were similar in the two groups. However, in OCD patients network borders extended further outwards from their centroids in each cortical zone. For example, in the anterior cingulate cortex, network borders shifted more anteriorly into the pregenual cortex, and, in lateral prefrontal cortex, network borders shifted more anteriorly towards the frontal pole (**Fig. 2d**).

### Contraction of frontostriatal executive networks

Given the prominent and robust frontostriatal expansion of the salience network in individuals with OCD, we asked the question whether such expansion affected cortical and striatal representation of other functional neighbouring networks. We focused our investigation on a subset of networks, namely the default (parietal), cingulo-opercular, and the frontoparietal network (**Fig. 3a**), which neighbour the salience network and are generally implicated in high level cognitive processes usually compromised in OCD patients. Expansion of the salience network in cortex was accompanied by contraction of the frontoparietal network in OCD individuals (**Fig. 3b**). On average, in OCD individuals the frontoparietal network occupied 28% less of the cortical surface relative to the mean in HC (8.73% ± 1.84% of cortex in OCD vs 12.09% ± 1.53% of cortex in HC) (**Fig. 3c**), with a large effect size (Cohen’s *d* = −2.03). This effect remained statistically significant when controlling for potential confounding effects such as sex and head motion (**Extended Data Fig. 5**) and with or without correction for potential site- or scanner-induced biases (**Extended Data Fig. 3c**). We also evaluated if representation of the frontoparietal network was similarly reduced in the striatum, given known functional and anatomical connections, but found that the difference in group means was not statistically significant. Instead, we observed a striatal reduction of the cingulo-opercular network in individuals with OCD (OCD: 20.47% ± 8.39 %; HC: 42.83% ± 13.52%; Cohen’s *d* = −1.89). The effect was consistently present when controlling for potential confounding factors such as sex, individual differences in head motion, age (**Extended Data Fig. 6**), and different acquisition sites (**Extended Data Fig. 3d**).

**Fig. 3.**
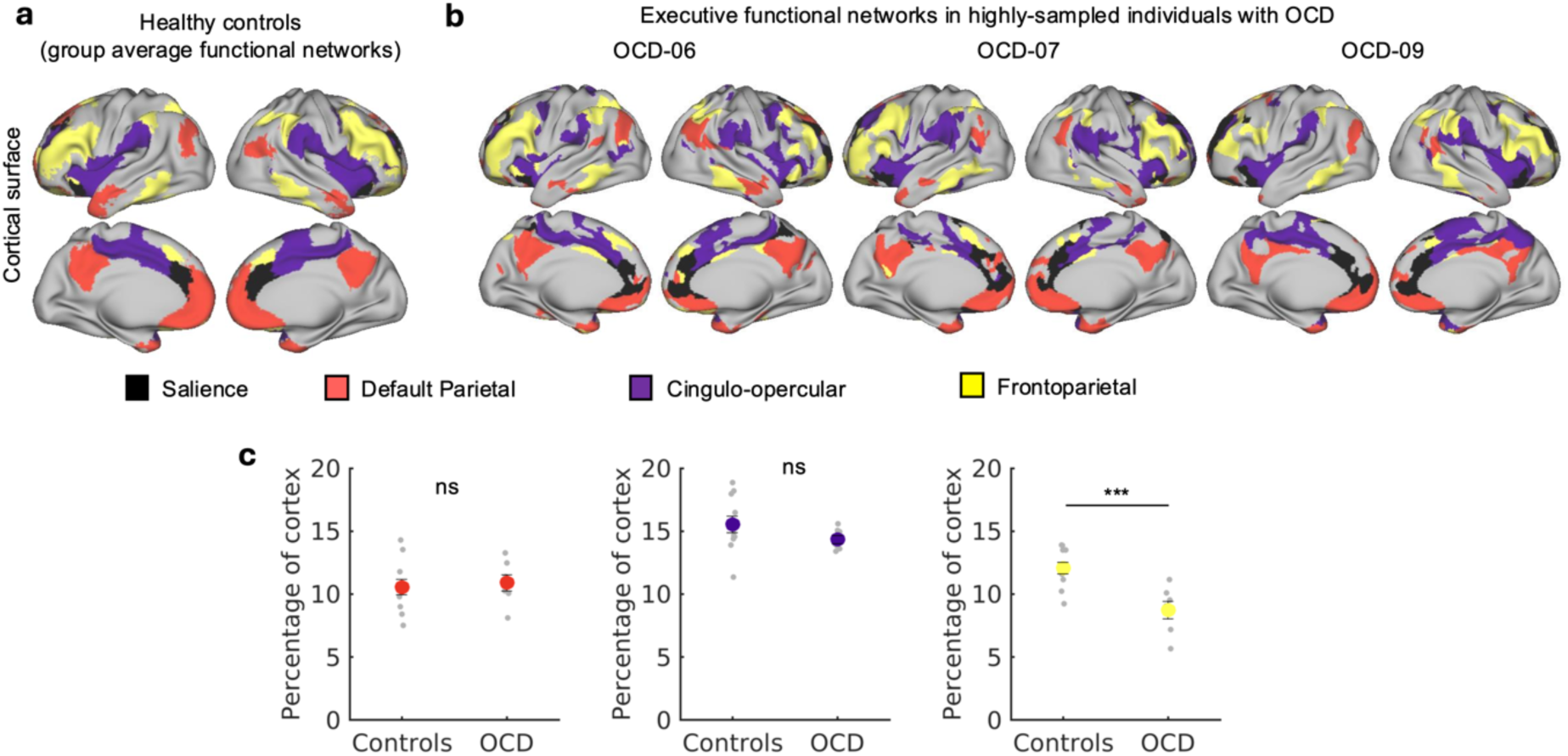
Expansion of the salience network is accompanied by frontostriatal contraction of neighbouring executive networks. **a,** Executive networks are shown in a group-average map of HC and **b,** in three representative subjects with OCD. Each of the executive networks investigated is shown with a different color (parietal subnetwork of the DMN, red; FP, yellow; CO, purple). **c,** Expansion of the salience network was accompanied by cortical contraction of the frontoparietal network in individuals with OCD (non-parametric Wilcoxon rank-sum *t*-test *P* = 0.001, rank sum = 32, Bonferroni corrected; Cohen’s *d* = −2.03) and striatal contraction of the cingulo-opercular network (non-parametric Wilcoxon rank-sum *t*-test *P* = 0.004, rank sum = 36, Bonferroni corrected; Cohen’s *d* = −2.03). Data are presented as mean ± s.d. DMN, default mode network; FP, frontoparietal; CO, cingulo-opercular.

### Salience network expansion affects heteromodal association areas

The observation that individuals with OCD exhibited cortical expansion in the salience network and concomitant contraction in the frontoparietal network raised the question of whether these phenomena were interrelated. Specifically, we explored whether the salience network’s expansion encroached upon neighbouring networks, particularly affecting heteromodal association areas like the frontoparietal network. Thus, to investigate the mechanisms giving rise to frontostriatal networks imbalances in OCD, we first generated a central tendency functional network map for HC (see Methods) (**Fig. 4a**). Next, we identified regions within the salience network of OCD individuals that either overlapped or did not overlap with the salience network in the group-average map of HC (**Figs. 4b, 4c**). For each individual with OCD, regions in the salience network that did not overlap with the HC group-average map were classified as “encroaching” (**Fig. 4c**, top), following the approach from^24^. For each subject, we quantified the degree of encroachment on other networks by calculating the relative contribution of each network to the total surface area of the encroaching salience network regions (**Fig. 4c**, bottom). This analysis revealed that salience network expansion was not random but predominantly targeted three neighbouring higher-order functional systems: the default mode (parietal and dorsolateral subnetworks), frontoparietal, and cingulo-opercular networks (**Fig. 4d**). These findings suggest that cortical border shifts drive frontostriatal imbalances in OCD, with expansion of the salience network resulting in encroachment on adjacent networks implicated in executive functions. Such shifts may reflect mechanisms governing cortical expansion, which refine functional boundaries during development or in response to environmental influences. To have a better understanding on the anatomical brain regions that were taken over by the salience network in OCD patients, we generated a group-average functional networks map for OCD patients. Next, we identified those regions in the salience network of the OCD group-average maps that were encroaching with respect to the group-average map generated for HC and labelled them based on a multi-modal parcellation of the human cerebral cortex^33^. We found that the salience network encroached on several areas including both medial and lateral regions of the orbitofrontal complex (OFC, Areas 11l, 11m), the ventrolateral prefrontal cortex (PFC) (Areas 47l, 47m) and a central region of the dorsolateral PFC (Areas 46, 9-46d, 9-46v) which are known to be characterized by abnormal functioning in OCD (**Supplementary Table 1** for a comprehensive list of regions).

**Fig. 4.**
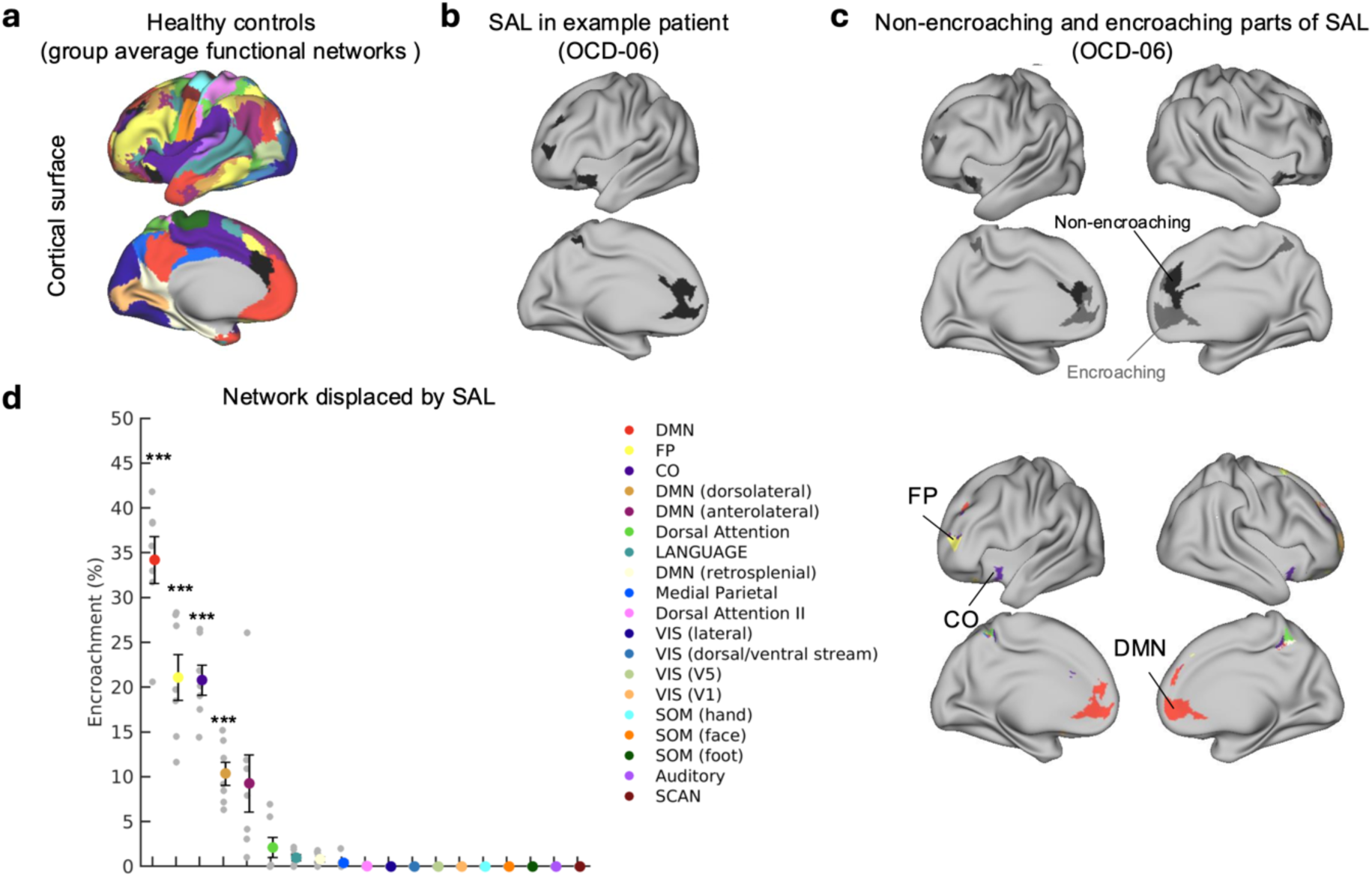
Encroaching of the salience network upon heteromodal, executive networks. **a,** Mode functional brain network assignments in cortex in HC. **b,** Salience network in a representative individual with OCD. **c,** Regions of the salience network of each individual with OCD that did and did not overlap with the HC average map are referred to as non-encroaching and encroaching, respectively. **d,** Salience network expansion selectively affected heteromodal networks, encroaching upon the DMN (parietal and dorsolateral subnetworks), frontoparietal and cingulo-opercular/action mode network. Statistical significance was assessed using two-tailed one-sample *t*-tests; all *P* < 0.001, Bonferroni corrected. Data are presented as mean ± s.d. CO, cingulo-opercular/action node; DMN, default mode; FP, frontoparietal.

### Frontostriatal connectivity tracks severity of OCD symptoms

The results above indicated individual-level specific topological features of the salience network in OCD patients. We then asked whether functional connectivity in frontostriatal circuits tracked symptom fluctuations, a hypothesis generated based on human^34,35^, and animal studies^36–38^ showing that the activity in frontostriatal circuits controls compulsive behaviours. Our analyses took advantage of the dense sampling longitudinal design in which each subject was tested multiple times over a significant period of time, spanning 3 months. This afforded an opportunity to ask how variability in brain network functional connectivity relates to fluctuations in specific symptom domains. Over the course of our dense-sampling longitudinal study, we observed significant fluctuations in different symptom dimensions, which were derived from standardized symptom scales and clinical interview. Interestingly, while in some individuals there were concomitant fluctuations in OCD and depression severity, in other individuals, the two trajectories were uncorrelated, possibly also due to the presence of comorbidities in some patients but not in others (**Fig. 5b**).

**Fig. 5.**
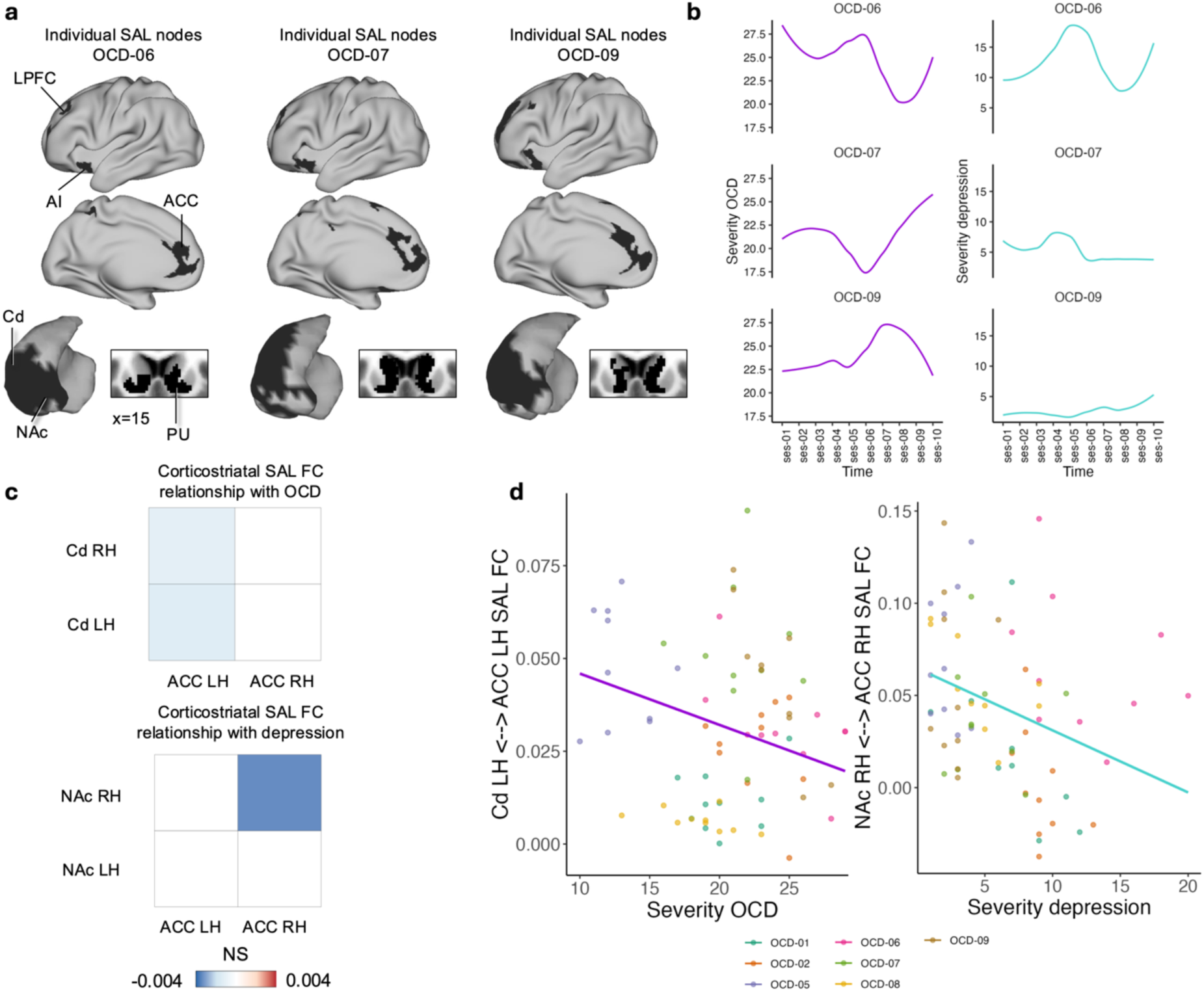
Frontostriatal salience network connectivity relates to fluctuations in OCD and depression severity over time in deeply sampled individuals with OCD. **a,** Frontostriatal nodes of the salience network in OCD-06, OCD-07, OCD-09. **b,** Fluctuations in OCD and depression severity from clinical interviews and self-report scales in deeply sampled individuals with OCD. **c,** FC between the salience network nodes in the Cd and the left ACC most closely tracked fluctuations in the severity of OCD symptoms as measured via the Y-BOCS (β = −0.001, *T* = −2.7, df = 29, *P* = 0.011, Bonferroni corrected, statistical significance was assessed using two-tailed permutation tests). FC between the right NAc and the right ACC most closely tracked fluctuations in the severity of depression symptoms (β = −0.003, *T* = −2.305, df = 50, *P* = 0.025, statistical significance was assessed using two-tailed permutation tests, but the result did not survive correction for multiple comparisons). In the figure, fixed-effect estimates for OCD symptom severity or depression that did not exceed chance (based on the null distribution) were set to zero. Only significant associations after permutation testing are shown. The predicted values from the model together with raw data are plotted in panel **d.** LPFC, lateral Prefrontal Cortex; AI, Anterior Insula; ACC, Anterior Cingulate Cortex; Cd, Caudate; NAc, Nucleus Accumbens; PU, Putamen; FC, Functional Connectivity. LH; Left Hemisphere; RH; Right Hemisphere.

To assess whether functional connectivity between specific salience network nodes tracked symptom fluctuations, we first identified nodes of the salience network in each individual subject based on individual specific topological features. (**Fig. 5a**). Next, we looked at the functional connectivity between a priori selected nodes and investigated the relationship between changes in functional connectivity strength over time and fluctuations in severity of OCD symptoms. As a cortical node, we focused on anterior cingulate cortex (ACC) salience node. ACC has been implicated in OCD pathophysiology^1^ and it encroached onto brain regions critically important in OCD (**Supplementary Table 1)**, such as for example the orbitofrontal complex which has been consistently found hyperactive at rest and during symptom provocation in fMRI studies in OCD^39,40^. Animal research has also demonstrated a causal role for the orbitofrontal cortex in modulating compulsive behaviour and habits^37^. We investigated its connectivity with salience nodes in the caudate (Cd) given recent evidence of optimal clinical improvements in OCD upon DBS of the anterior limb of the internal capsule extending to the caudate nucleus^35^, also in line with animal studies^41^. Overall, these areas within the frontostriatal circuits have been shown to regulate symptom severity in OCD in several neuroimaging studies^34^. We found that the connectivity between the ACC and the Cd (dorsal striatum) nodes of the salience network was negatively correlated with changes in OCD symptom severity (**Figs. 5c, 5d**). This analysis included head motion as a covariate and the significance of the effect was confirmed via permutation testing. Given previous reports of a relationship between the ACC node of the salience network and the nucleus accumbens with symptom severity in individuals with depression^24^, we also tested the relationship between the functional connectivity between these nodes and depression. We found that the ACC - NAc connectivity negatively correlated with severity of depression in an analysis including head motion as a covariate and whose significance was confirmed via permutation testing (**Figs. 5c, 5d**), though it did not survive correction for multiple comparisons. We also conducted a specific analysis testing specifically the association between ACC-NAc connectivity and anhedonia in line with previous reports^24^. We identified only those items related to anhedonia based on clinical decision and conducted an analysis to investigate the relationship between ACC-NAc and anhedonia but we failed to find a significant association. However, it must be noted that we only had a limited subset of items related to anhedonia (**Supplementary Table 2**). Additionally, our sample included only OCD patients, and most of them were free of comorbidities. Hence, it is possible that variations in anhedonia might not be pronounced.

## Discussion

Our findings from this precision imaging study in densely sampled individuals with OCD showed that there is a striking imbalance within frontostriatal networks. Cortical and striatal expansion of the salience network was concomitant to marked contraction of the frontoparietal network. These alterations were robust to methodological variations and exhibited large effect sizes, standing in stark contrast to the more subtle effects typically reported in conventional neuroimaging studies. The expansion of the salience network was driven by spatially organized border shifts that spared unimodal sensorimotor areas and selectively encroached on functional networks which are implicated in high-order cognitive functions. From a symptomatology standpoint, such changes are particularly intriguing, as OCD individuals appear to have an impaired ability to correctly modulate the emotional significance of internal and external stimuli—a function typically ascribed to the salience network^42^. Similarly, the frontoparietal network is typically associated to goal-directed behaviour abilities. Its contraction aligns with the characteristic behavioural imbalance observed in OCD patients, who often exhibit impaired goal-directed control^43–45^ and, more broadly, executive dysfunction^46–48^ (see also, Chapter 22 in^49^) making them increasingly reliant on habitual behaviours. Together these correlated network expansion-contraction dynamics provide a strong neurobiological substrate for several key behavioural traits of OCD, in line with the idea that such networks reconfiguration might underpin changes in key pathological behavioural traits typically observed in OCD. In addition to evaluating network topologies with individual precision, we availed of longitudinal data to investigate the relationship between changes in functional connectivity and fluctuations in symptom severity. Our findings showed that dynamic changes in striatal connectivity with the anterior cingulate node of the salience network tracked OCD symptom severity.

An outstanding question pertains to the exact mechanisms driving network expansions and contractions in OCD. The existing literature suggests that both genetic factors^50,51^ and activity-dependent processes may contribute to individual differences in network topology. For example, network topology can be affected by motor training which can expand M1 representation, while disuse (due to amputation, casting, or congenital defects) reduces it, allowing other body parts to take over^52,53^. Similarly, area V1 in congenitally blind individuals is significantly decreased in size^54–59^. Such foundational mechanisms are conserved across species^60^. While the majority of evidence pertains to low level motor and visual systems, recent work has shown that similar mechanisms extend to high level processing brain regions in humans. Long-term working memory training altered the role of the prefrontal cortex, which developed representations for specific stimuli and associations learned over time^61^. Likewise total cortical representation of the frontoparietal network was found to be positively correlated with executive function abilities in children^62^ and individual network topologies in association systems related to different neuropsychological profiles^14^. In OCD, an expansion of the salience network alongside a contraction of the frontoparietal network may represent a reallocation of cortical territory driven by functional demands. Patterns of hyper- and hypo-activation generally observed in MRI studies of OCD might relate to imbalanced network topologies observed here. The salience network has critical nodes in the ACC and in OCD individuals it encroached on areas such as the orbitofrontal cortex. Both of these brain areas are part of a longstanding neuroanatomical model of OCD that centres on frontostriatal circuits^1,63,64^. Hyperactivity of the orbitofrontal cortex is one of the most replicated neuroimaging findings in OCD. Increased activation has been found at rest and during symptom provocation and it is related to pharmacological response to SSRIs^65,66^. Additionally, cingulotomy has been successful for otherwise treatment resistant OCD^67,68^ and, more recently, a neurochemical imbalance between glutamate and GABA in the ACC has been linked to clinical compulsive symptoms^69^.

In contrast, a pattern of hypoactivation and reduced connectivity is found in associative cortices during tasks requiring top-down control in OCD patients^46,47,70,71^. These results together with the observation of compromised executive control^48,72,73^ can be linked to the marked contraction of frontoparietal network observed for the first time here. Overall, the observed imbalance appears to create a foundation for disrupted regulation of bottom-up emotional states and impaired top-down control of goal-directed behaviours. Future work will enable the investigation of whether individual network topologies have consequences for functional and behavioural variations during cognitive tasks and if these might represent candidate endophenotypes for example through the investigation of network topology in first-degree unaffected relatives.

The discovery of individual-specific network topologies in OCD has significant implications for designing therapeutic neuromodulation interventions, which may have widely varying effects due to inter-individual differences. For instance, both transcranial magnetic stimulation^74^ and deep brain stimulation^35,75^ are promising treatment avenues for OCD. However, the presence of distinct network organizations at the same anatomical location across individuals poses a challenge for neuromodulatory approaches—not only aimed at modulating activity of the striatum (as in deep brain stimulation) but also those targeted at manipulation of cortical brain circuits and networks^76^. Group-average parcellations may not accurately reflect an individual’s brain organization, contributing to signal mixing and reducing targeting precision, leading to mixed effects or reduced efficacy for a given patient. Previous research has shown that the frontoparietal network is among the most individually variable^14,77^, and our findings suggest that this network is particularly susceptible to expansion mechanisms of the salience network in OCD. Given the idiosyncratic functional organization of OCD, it is highly likely that standard neuromodulatory approaches engage different functional networks in different individuals. It is also plausible that lack of response in some patients may be due in part to failure to engage the desired functional network. Therefore, insights from the current study could pave the way for more personalized interventions in OCD. For example, transcranial magnetic stimulation protocols that account for a patient’s unique functional network topology have shown promise in depression^78^, and our findings provide a strong rationale for applying a similar individualized approach in OCD.

Expansion of the salience network has been observed in patients with depression^24^. Our finding of a similar expansion in patients with OCD challenges the specificity of salience network enlargement as a biomarker for depression. Instead, these results suggest that salience network vulnerability, leading to variations in network topology, may serve as a transdiagnostic marker of susceptibility to psychiatric disorders in general. This is also in line with the observation that, in the previous study^24^, most individuals with depression also had comorbid diagnoses, including OCD. Regardless of this, the large effect sizes associated with expansion of the salience network suggest that it might serve in the future as a metric for assessment and treatment of debilitating psychiatric conditions. An additional hypothesis is that the specific form of psychopathology an individual develops may be influenced by network topology variations associated with expansion of the salience network. For instance, in this study, we found that the salience network expanded into regions such as the orbitofrontal cortex. Moreover, this expansion was accompanied by a contraction of the frontoparietal network—a finding that, to the best of our knowledge, is unique to OCD and has not been identified in previous reports across multiple independent samples of individuals with depression^24^. This might suggest that the specific pattern of salience network expansion alongside frontoparietal contraction may represent a distinct neural phenotype associated with OCD. Studies with bigger sample sizes will also enable to address specifically the role of comorbidities. Finally, previous work indicated that the salience network was not larger than normal in individuals with OCD^24^. However, such results were derived from a dataset with only 1 hour of multi-echo fMRI data per-subject. By using that amount of data in our sample we also failed to identify a difference between OCD individuals and HC (**Extended Data Fig. 7**). Therefore, this analysis can explain why previous work could not find expansion of the salience network in OCD individuals ^24^. Previous work also highlighted that precise and reliable mapping of the salience network and consistent detection of its expansion may require 1.5-2h of high-quality fMRI data per subject^24^. Overall, from a methodological point of view, our analyses confirmed that reliable mapping of individual differences in network topology requires the use of precision functional mapping in combination with large quantities of high-quality, densely sampled multi-echo fMRI data per-subject.

While this might mean that extensive imaging is needed in patients, functional scans are relatively non-invasive and the approach required by PFM is not substantially more demanding than comprehensive structural MRI protocols currently in use, suggesting this a promising tool for clinical translation. Capitalizing on the longitudinal design and the densely sampled experimental paradigm, we investigated whether changes in functional connectivity between nodes of the salience networks related to fluctuations in symptoms. Results showed a dissociation whereby the caudate tracked to severity of OCD, while a ventral network implicating the nucleus accumbens was associated with symptoms of depression. The latter did not survive correction for multiple comparisons, but the results are in line with previous reports^24^. Critically, recent study aimed at identifying optimal stimulation sites for OCD-DBS has found a sweet spot in the anterior limb of the internal capsule extending to the caudate nucleus^35^. In contrast, the nucleus accumbens proper was associated with beneficial but suboptimal clinical improvements. Overall, these findings move us closer towards a circuit-based understanding for OCD. Extensive longitudinal imaging in single subjects will enable to investigate these associations per subject, possibly guiding tailored intervention.

To conclude, our study provides proof-of-principle data to support the use of precision functional mapping and deep, longitudinal sampling for understanding circuits at the subject level in psychiatry. Our analyses show a frontostriatal imbalance in OCD in the form of an expansion and contraction of the salience and frontoparietal network, respectively. Additionally, the use of precision functional mapping allowed delineation of functional networks at the individual patient level with implications for tailored intervention. Finally, the use of extended data acquisition showed that reduced connectivity in a dorsal frontostriatal circuit tracked severity of obsessions and compulsions, opening new avenues to the understanding of neurobiological mechanisms implicated in the emergence and remission of symptoms in OCD.

## Methods

### Datasets

Data were collected from 7 individuals (median age 30, range: 22-48; 3 female (F)/ 4 male (M)) with a diagnosis of OCD (based on DSM-IV-TR criteria and confirmed by the Mini-International Neuropsychiatric Interview). At the time of testing, most (5/7, 71%) individuals with OCD in our study sample did not have any comorbidity. The remaining part of the sample (2/7, 29%) had a comorbid diagnosis (major depressive disorder [MDD] in remission, *n* = 1; MDD and Alcohol Use Disorder, *n* = 1) though OCD remained the chief complaint. With respect to medication status, most (4/7, 57%) individuals were unmedicated being either drug-naïve or off medication for more than 8 weeks prior taking part of the study. Medicated (3/7; 43%) patients were taking SSRI (*n* = 1), SNRI (*n* = 1), or SSRI in combination with antiepileptic drugs (*n* = 1). The sample size was based on funding availability and timing; no statistical power analyses were performed. However, the number of individuals tested in this study is in line with previous reports of highly sampled clinical and non-clinical populations^12,22,24,61,79^.

To compare our findings against the HC group of densely-sampled individuals, we included data from two sources – the MyConnectome (*n =* 1, 45-year-old male)^10,11^ and the Midnight Scan Club (MSC; *n* = 10, mean age 29.1± 3.3 years, 5F/FM)^12^. Data from the MyConnectome and the MSC were obtained online (https://openneuro.org/) in a preprocessed, fully denoised and surface-registered format, and no further preprocessing or denoising was performed for the present study.

### Study Design

For each individual with OCD, data acquisition was performed over the course of 10 sessions, conducted on separate days. Two participants had an extra session due to excessive head motion or reported sleepiness over the course of previous scanning sessions which were not retained for data analysis. Sessions were distributed throughout the study, so that each participant had a scan weekly. All sessions for all participants took place at the Stanford Center for Cognitive and Neurobiological Imaging, Stanford University. To monitor fluctuations in specific symptoms domain longitudinally, on each experiment day, participants were administered the Y-BOCS^32^. To assess symptoms of depression participants were also asked to complete the Depression and Anxiety and Stress Scale (DASS-21)^80^ clinical scale. Next, they underwent MRI acquisition entailing two behavioural tasks and resting-state fMRI. The sequence of events inside the scanner was maintained fixed across subjects and sessions. Finally, in one of the 10 functional sessions, structural data were acquired at the end of that session. The majority of scans for each participant took place approximately at the same time of the day, limiting significant AM-PM fluctuations^81^. Accordingly, for all participants scanning on separate days was conducted with maximum 3 hr lag, with the exception of two scanning sessions in two participants, and one scanning session in other two participants. In each of the individuals with OCD, we collected 3 hr of resting-state fMRI data and 2.8 hr task-based fMRI data across two different behavioural tasks. Analysis of task-based data is ongoing and not reported here. The study was approved by the Stanford Institutional Review Board (Protocol number: 63831). Participants gave informed consent prior to participation.

### MRI acquisition

Data were acquired on a GE Signa 3T MRI Scanner at the Center for Cognitive and Neurobiological Imaging on Stanford University campus using a Nova Medical 32-channel head coil. In each session, resting state fMRI was acquired over the course of three separate 6-min runs as previous studies found that sampling over shorter epochs increased reliability^11^. Accordingly, three multiecho, multi-band resting-state fMRI scans were collected using a T2*-weighted echo planar sequence covering the full brain (TR 1,490 ms; TE1 13.4 ms, TE2 36.4 ms, TE3 59.4 ms; flip angle 53°; 2.8 mm isotropic voxels; 51 slices; AP phase encoding direction; and multi-band acceleration factor 3) with 242 volumes acquired per scan for a total acquisition time of 6 min. In each session, a dual-echo spiral field map sequence was acquired with the same prescription as the functional images. T1-weigthed (TR 3000 ms; TE 3.5 ms; flip angle 8° and 230 sagittal slices with 0.8 mm slice thickness) and T2-weighted anatomical images (TR 2825 ms; TE 96 ms and 186 sagittal slices with 0.9 slice thickness) were acquired for each subject. Structural data were always collected at the end of the dedicated session (i.e., after the functional scans). Anatomical and functional data were preprocessed using *fMRIPrep* 22.0.0rc4^82^ as described below.

### Anatomical preprocessing and cortical surface generation

Anatomical data (T1-weighted image, T1w) were corrected for intensity non-uniformity (INU) with N4BiasFieldCorrection^83^, distributed with ANTs 2.3.3^84^, and used as T1w-reference throughout the workflow. The T1w-reference was then skull-stripped with a *Nipype* implementation of the antsBrainExtraction.sh workflow (from ANTs), using OASIS30ANTs as target template. Brain tissue segmentation of cerebrospinal fluid (CSF), white-matter (WM) and gray-matter (GM) was performed on the brain-extracted T1w using fast (FSL 6.0.5.1^85^). Brain surfaces were reconstructed using recon-all (FreeSurfer 7.2.0^86^), and the brain mask estimated previously was refined with a custom variation of the method to reconcile ANTs-derived and FreeSurfer-derived segmentations of the cortical gray-matter of Mindboggle^87^. Volume-based spatial normalization to one standard space was performed through nonlinear registration with antsRegistration (ANTs 2.3.3), using brain-extracted versions of both T1w reference and the T1w template. The following template was selected for spatial normalization: *ICBM 152 Nonlinear Asymmetrical template version 2009c* (MNI152NLin2009cAsym).

### fMRI preprocessing

Functional multiecho preprocessing of resting-state data entailed the following steps. First, a reference volume and its skull-stripped version were generated from the shortest echo of the BOLD run using a custom methodology of *fMRIPrep*. Head-motion parameters with respect to the BOLD reference (transformation matrices, and six corresponding rotation and translation parameters) are estimated before any spatiotemporal filtering using mcflirt (FSL 6.0.5.1^88^). The estimated *fieldmap* was then aligned with rigid-registration to the target EPI (echo-planar imaging) reference run. The field coefficients were mapped on to the reference EPI using the transform. BOLD runs were slice-time corrected to 0.701s (0.5 of slice acquisition range 0 s-1.4 s) using 3dTshift from AFNI^89^.

The BOLD reference was then co-registered to the T1w reference using bbregister (FreeSurfer) which implements boundary-based registration^90^. Co-registration was configured with six degrees of freedom. First, a reference volume and its skull-stripped version were generated using a custom methodology of *fMRIPrep*. Several confounding time-series were calculated based on the preprocessed *BOLD* including framewise displacement (FD) and three region-wise global signals. FD was calculated for each functional run using its implementations in *Nipype* (following the FD definition by^91^). The three global signals are extracted within the CSF, the WM, and the whole-brain masks. The head-motion estimates calculated in the correction step were also placed within the corresponding confounds file. The confound time series derived from head motion estimates and global signals were expanded with the inclusion of temporal derivatives and quadratic terms for each^92^. The distortion-corrected (STC, HMC, SDC) per echo timeseries were then used independently with multi-echo tools to perform more advanced denoising as detailed below.

### Multi-echo denoising

Individual preprocessed data for all echoes were submitted to multi-echo ICA (ME-ICA^30^), which is designed to isolate spatially structured T2*-dependent (neurobiological; BOLD-like) and S0-dependent (non-neurobiological; non BOLD-like) signals and implemented using the tedana.py workflow^93^. In short, the preprocessed, ACPC-aligned echoes were first combined according to the average rate of T2* decay at each voxel across all time points by fitting the monoexponential decay, *S*(*t*) = S0e^−*t*/T2*^. From these T2* values, an optimally combined multi-echo (OC-ME) time series was obtained by combining echoes using a weighted average (WTE = TE × e−^TE/T2*^). The covariance structure of all voxel time courses was used to identify main signals in the OC-ME time series using principal component and independent component analysis. Components were classified as either T2*-dependent (and retained) or S0-dependent (and discarded), primarily according to their decay properties across echoes. All component classifications were reviewed by M.M.V.

### Surface processing and CIFTI generation of fMRI data

The denoised fMRI time series were mapped to the individual’s fsLR 32k midthickness surfaces with native cortical geometry preserved (using the -ribbon-constrained method), combined into the connectivity informatics technology initiative (CIFTI) format^94^. This was achieved via Ciftify’s ciftify_subject_fmri pipeline. This yielded time courses representative of the entire cortical surface, subcortex (accumbens, amygdala, caudate, hippocampus, pallidum, putamen, thalamus, cerebellum, diencephalon, and brainstem) and cerebellum but excluding non-grey matter tissue. After preprocessing and ME-ICA denoising, time-series data were further processed by regressing out confounds including motion, white matter, csf, and global signal and applying a temporal Butterworth bandpass filter (0.1 – 0.01) using *Nilearn*. Temporal masks were generated for censoring high-motion time points. Motion-contaminated volumes were identified by framewise displacement (FD) as quantified through fMRIPrep using the formula proposed by^91,95^. Frames with FD > 0.3 mm were flagged as motion-contaminated and censored. Across all subjects, these masks censored 3% ± 3% (range: 0% - 7%) of the data; on average, subjects retained 6872 ± 184 volumes (range: 6582 – 7050). Note that in this study, even the worst subject retained more than 2.5 hours of data. Spurious coupling between subcortical voxels and adjacent cortical tissue was mitigated by regressing the average time series of cortical tissue of less than 20 mm in Euclidean space from a subcortical voxel. Finally, time-series were spatially smoothed with geodesic (for surface data) and Euclidean (for volumetric data) Gaussian kernels (*σ* = 2.55 mm) using Connectome Workbench command line utilities^83^.

### Evaluating the role of individual identity to network similarity

We first aimed to establish the contribution of group-level and individual-specific effects to network similarity. To this end, to evaluate the within- and across-subject similarity, we first generated a parcel-to-parcel functional connectivity matrix for each subject in each session. We used a common set of cortical parcels^33^ and computed FC by averaging the preprocessed and denoised BOLD time course within each parcel. FC was computed by averaging the preprocessed and denoised BOLD time course within each parcel. FC values were Fisher transformed for normality. Accordingly, for each subject and session FC was represented with a parcel × parcel functional network matrix. Firstly, we used a classical multidimensional scaling (MDS) approach to depict how individual variance affected the similarity among networks in a data driven fashion and quantified the variance explained with this approach with principal component analysis (PCA). MDS places data in multidimensional space based on the similarity (Euclidean distance) among data points – where in this case a data point represents the linearized upper triangle of a given functional network matrix. Each separate matrix (from a given subject, and session) was entered into the classical MDS algorithm (implemented in Python with Scikit-Learn, MDS). Then, the effects of group and individuals were directly examined by calculating the similarity among each original functional network matrix (i.e., correlation among the linearized upper triangles), creating a second-order ‘‘similarity’’ matrix, following the same approach adopted in previous studies^17^. Subsequently, the average similarity was examined for functional network matrices that were from different individuals, tasks, and sessions (i.e., group effect) and from the same subject by different sessions (i.e., individual effect). These effects were calculated per subject and were compared with one another using paired two-tailed t tests. These procedures collectively correspond to the analyses performed in the section “Individual identity dominates functional networks” and **Fig. 1**.

### Precision functional mapping of functional brain networks in individuals

Following removal of motion-contaminated volumes as described above, data analysis was based on an average of 170.66 min of multi-echo resting-state fMRI scanning for OCD subjects (range: 163.45-175.08 min) across 10 sessions and an average of 271.88 min (range: 100-816.66 min) for HC (range sessions:10-84). Therefore, amount of data included for each of our OCD and HC individuals is above what previous work has shown to be necessary for reliable network mapping (i.e., 90 to 120 minutes)^24^. We computed a functional connectivity matrix for each individual quantifying the correlation between the time courses of all cortical vertices and subcortical voxels, concatenated across all study visits. Geodesic and Euclidean space were used for cortico-cortical and subcortical-cortical distance respectively to set correlations between nodes 10 mm or less apart to zero, to avoid basing network membership on correlations attributable to spatial smoothing. Correlations between voxels belonging to subcortical structures were set to zero. To delineate functional brain networks and their boundaries at the individual level we used the InfoMap community detection algorithm^96^ which is a widely used approach^10–12,24,97^. The algorithm was run at different density thresholds so that the strongest X% correlations to each vertex and voxel were retained (0.01%, 0.02%, 0.05%, 0.1%, 0.2%, 0.5%, 1%, 2% and 5%). Free parameters such as the number of algorithm repetitions (i.e., 50) for the InfoMap algorithm were fixed across subjects. The level of connectivity retained in the functional connectivity matrix following thresholding affects the number of communities identified by the InfoMap algorithm. Following previous implementations of this procedure^24^, we selected 0.1% graph density as the optimal scale for further analyses. This resulted in 96.67 ± 16.93 communities on average across individuals which is in line with previous reports^24^. Functional connectivity priors were used to algorithmically assign each InfoMap community to one of 20 possible functional networks identities (Default-Parietal, Default-Anterolateral, Default-Dorsolateral, Default-Retrosplenial, Visual-Lateral, Visual-Stream, Visual-V1, Visual-V5, Frontoparietal, Dorsal Attention, Premotor/Dorsal Attention II, Language, Salience, Cingulo-opercular/Action-mode, Parietal memory, Auditory, Somatomotor-Hand, Somatomotor-Face, Somatomotor-Foot, Auditory or Somato-Cognitive-Action). Accordingly, network identities were assigned based on functional connectivity and spatial locations of each community relative to the specified set of priors. A confidence score is assigned to each assignment and computed as the relative difference in total score associated with the winning and the first runner-up network assignment. Algorithmic assignments with a confidence value below 0.33 were visually inspected as recommended in previous work ^24^, but no manual adjustment was carried out.

### Quantification of functional network size and spatial locations in individuals

We measured the relative contribution (size) of each functional network in each individual, following a validated pipeline^24^. To this end, we first measured the surface area (in mm2) associated with each vertex in the individual’s midthickness surface using the *--surface-vertex-areas* function from the Workbench Command within the Connectome Workbench. Next, we computed the total surface area of all network vertices in relation to the total cortical surface areas. This approach controls for the fact that each cortical vertex represents a different amount of surface area. For subcortical networks in the striatum, in which each voxel represents the same amount of tissue, we computed the relative contribution of each functional network to the total striatal volume by taking the total number of network voxels in relation to the total number of striatal voxels. Non-parametric Wilcoxon rank-sum tests were used to assess statistical significance in group differences in network size using the Matlab ranksum.m function. Assumptions about equal variance were adjusted when appropriate (based on two-sample *F*-tests performed using Matlab vartest2.m function). Cohen’s *d* effect size was calculated as the difference in group means divided by pooled standard deviation. The relative difference between groups was calculated as the absolute difference divided by the network size in HC. Density maps were created by calculating the percentage of individuals with salience network representation at each cortical vertex using the *-cifti-label-probability* function from the Workbench Command. These procedures collectively correspond to the analyses performed in the sections “Expansion of frontostriatal salience network in individuals with OCD,” “Contraction of frontostriatal executive networks,” and **Figs. 2-3**.

### Organization of functional networks following expansion of the salience network

To identify regions of each OCD individual’s salience network map that did or did not overlap with salience network assignment in HC, we proceeded as follows. Firstly, we generated a group-average HC map by calculating the mode assignment across HC at each point in the brain. More specifically, each vertex was assigned a functional network identity corresponding to the most frequently observed network identity across HC. Vertices assigned to the salience network in both an OCD individual and in the HC mode assignment map were operationalized as “non-encroaching.” In contrast, vertices that were assigned to the salience network in a given OCD individual but not in the group-average HC map were operationalized as “encroaching”. For each subject, an encroachment profile, was calculated as the relative contribution of each functional network to the total surface area of the encroaching portion of the salience network. We also computed a group-average OCD map and made a comparison between the group-average map from HC and OCD individuals. Encroaching portions of the salience network in group-average OCD map were mapped onto a multi-modal parcellation of the human cerebral cortex atlas^33^ to identify the functional and anatomical identity of those vertices assigned to the salience network in OCD individuals, but not in HC. These procedures collectively correspond to the analyses performed in the section “Salience network expansion affects heteromodal association areas” and **Fig. 4**.

### Longitudinal analyses relating changes in frontostriatal functional connectivity with symptom severity

To assess OCD symptom severity, we interviewed OCD individuals using the Y-BOCS (the gold-standard measure for OCD symptoms) at every session. At each session we also asked subjects to complete a clinical scale measuring severity of depression. the Depression, Anxiety and Stress Scale (DASS-21) from which specific items related to emotional states of depression, anxiety and stress were obtained. In order to conduct our analyses, we retrieved items of the depression subscale and obtained a score for each subject at each session. To derive an anhedonia-related symptom measure, the clinician (C.I.R.) rated the extent to which each DASS-21 item reflected anhedonia, and a separate score was computed using only these items for a dedicated analysis. Functional connectivity strength between pairs of nodes was based on individual-specific nodes of the salience network based on individual topologies. Accordingly, in each individual, the ACC node for example corresponded to the cluster located in the ACC and assigned to the salience network. We *a priori* selected cortical (anterior cingulate cortex) and striatal (nucleus accumbens, caudate) nodes based on specific working hypotheses. To test whether longitudinal changes in functional connectivity were related to symptom severity fluctuations, we used Linear Mixed-Effects Models (LMMs) via the *lmer* function from *lmerTest* in R. We specified two separate models: one examining the relationship between functional connectivity and OCD symptom severity and another assessing the relationship between functional connectivity and depression severity. Each model also included head motion as a covariate. A first model including random slopes for the variables of interest (i.e., severity of OCD symptoms and severity of depression) resulted in either singular fits or failed to convergence. Hence models with just a random intercept were carried forward. Statistical significance was determined via permutation testing using *bootMer* from *lme4*, which performs bootstrapping for mixed models. The same procedure was followed for the analysis specific to anhedonia symptoms. These procedures collectively correspond to the analyses performed in the section “Frontostriatal connectivity tracks severity of OCD symptoms” and **Fig. 5**.

## Data availability

Data from the MyConnectome dataset are available on OpenNeuro repository at https://openneuro. org/datasets/ds000031/versions/2.0.2. Data from the MSC dataset are available in the OpenNeuro repository at https://openneuro.org/data-sets/ds000224/versions/1.0.4. Raw data from the 7 individuals with OCD will be made available upon request via a Data Use Agreement, due to the sensitive nature of these data.

## Code availability

Code for performing the precision functional mapping and network size calculations described in this manuscript are maintained in an online repository (https://github.com/MatildeVaghi/PFM). Software packages incorporated into the above pipelines for data analysis included: Matlab R2022b, https://www.mathworks.com/; Connectome Workbench 1.4.2, http://www.humanconnectome.org/software/connectome-workbench.html; Freesurfer v7, https://surfer.nmr.mgh.harvard.edu/; FSL 6.0, https://fsl.fmrib.ox.ac.uk/fsl/fslwiki; and Infomap v2.0.0, https://www.mapequation.org. Advanced Normalization Tools (ANTS; v2.3.3), Nilearn (https://nilearn.github.io/stable/index.html).

## Author Contributions

Conceptualization: M.M.V., C.I.R., R.A.P.; Methodology: M.M.V., R.A.P.; Data acquisition: M.M.V., S.S., J.A.H.R.; Data analysis: M.M.V., M.N., R.A.P.; Resources: C.I.R.; Writing – Review and Editing: M.M.V., M.N., S.S., J.A.H.R., P.G.B., C.I.R., R.A.P.; Funding acquisition: M.M.V., R.A.P.

## Acknowledgements

We thank the study participants for their time and efforts in participating in this study. This work was supported by grants to M.M.V. from the Human Frontier Science Program Organization (LT000751/2019-L) and the Brain & Behaviour Research Foundation (Young Investigator NARSAD Award, 28751). M.M.V. is also supported by an MRC Career Development Award (MR/Y011384/1). Martin Norgaard was supported by the BRAIN Initiative grant (OpenNeuroPET, grant ID 1R24MH120004-01A1).

---

**Extended Data Fig. 1.**
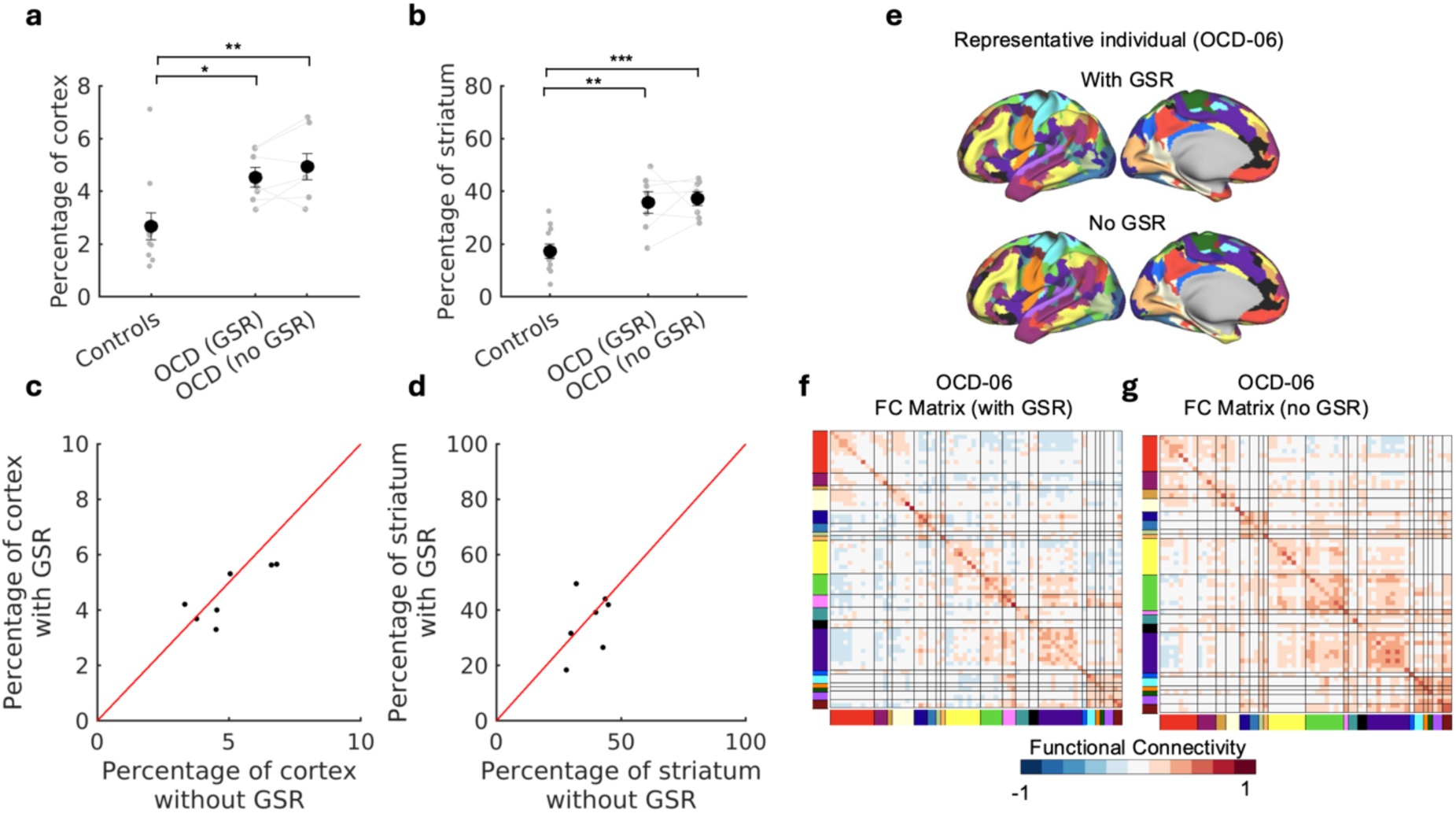
Effect of global regression. **a**, Group differences between OCD and healthy controls in cortical salience network expansion size remain statistically significant when Global Signal Regression (GSR) is not performed (non-parametric Wilcoxon rank-sum t-test, P=0.006, rank sum = 96, Cohen’s d = 1.45). **b,** Similarly, group differences between OCD and healthy controls in striatal salience network expansion remain statistically significant when GSR is not performed (non-parametric Wilcoxon rank-sum t-test, P<0.001, rank sum = 102, Cohen’s d = 2.44). No significant differences in cortical or striatal salience network expansion were observed in OCD patients, regardless of whether GSR was included in the analyses. **c,** Correlation of individual differences in cortical salience network size with (y-axis) and without (x-axis) GSR (Spearman correlation = 0.75). **d,** Correlation of individual differences in striatal salience network size with (y-axis) and without (x-axis) GSR (Spearman correlation = 0.46). **e**, All functional networks mapped on a single representative individual with OCD with and without GSR applied. **f, g**, Functional connectivity matrices for a representative individual with OCD with (**f**) and without (**g**) GSR. Collectively, these results indicate that GSR has little effect on functional brain network topography.

**Extended Data Fig. 2.**
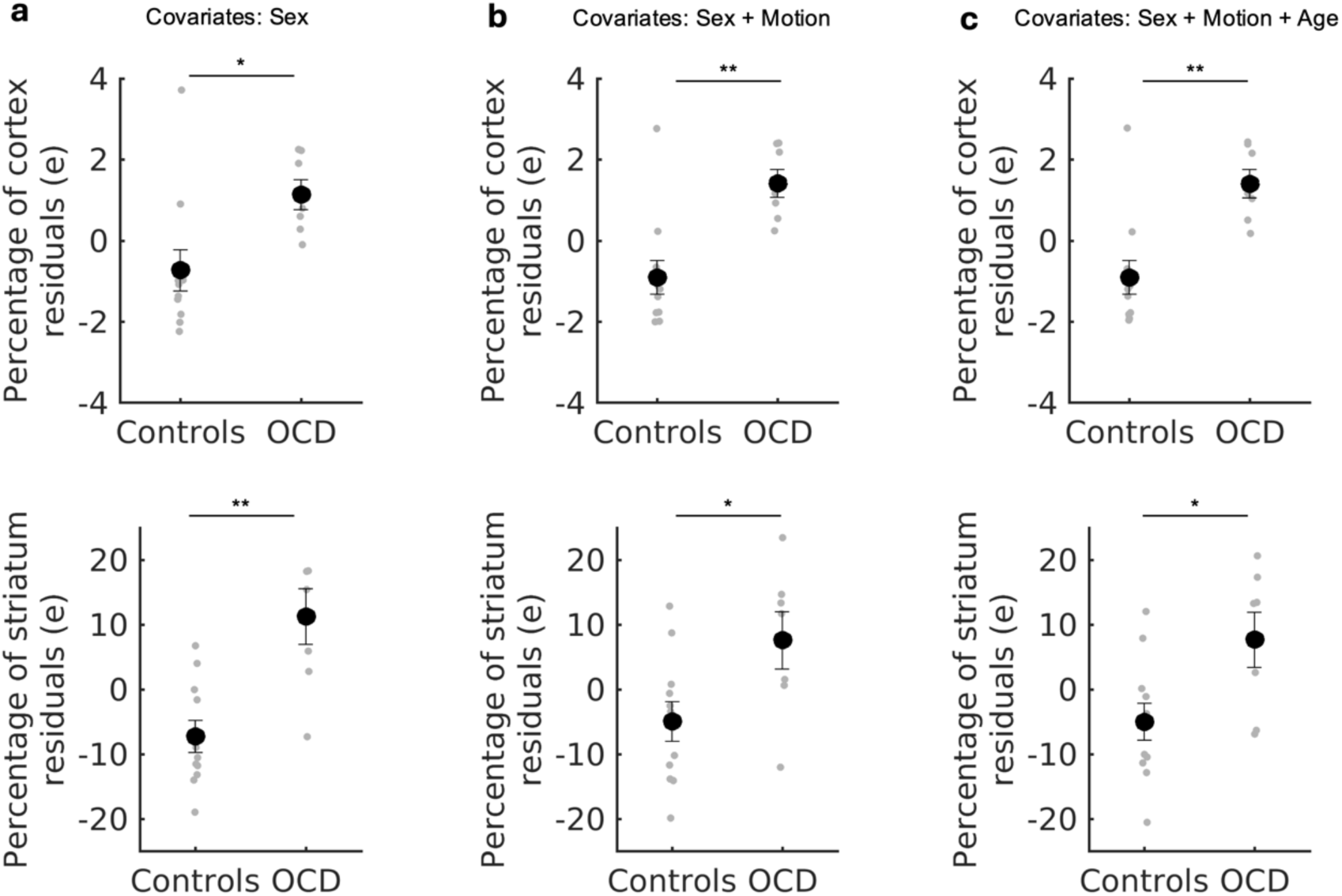
Cortical and striatal salience network expansion in OCD remain statistically significant when controlling for sex ratio and individual differences in head motion and age. **a,** The size of the salience network was regressed against sex, and group comparisons were repeated using the residuals (e). The salience network remained significantly larger in individuals with OCD compared to healthy controls, both in cortical (non-parametric Wilcoxon rank-sum t-test, P = 0.011, rank sum = 94) and striatal regions (non-parametric Wilcoxon rank-sum t-test, P = 0.003, rank sum = 98). **b-c**, This analysis was repeated with head motion (measured as the percentage of volume retained after motion censoring) and age (in years) as additional covariates. In all models, the salience network remained significantly larger in OCD individuals, both in the cortex (Sex + Motion: Wilcoxon rank-sum t-test, P = 0.003, rank sum = 98; Sex + Motion + Age: Wilcoxon rank-sum t-test, P = 0.004, rank sum = 97) and striatum (Sex + Motion: Wilcoxon rank-sum t-test, P = 0.027, rank sum = 91; Sex + Motion + Age: : Wilcoxon rank-sum t-test, P = 0.027, rank sum = 91). Data are presented as mean ± standard deviation.

**Extended Data Fig. 3.**
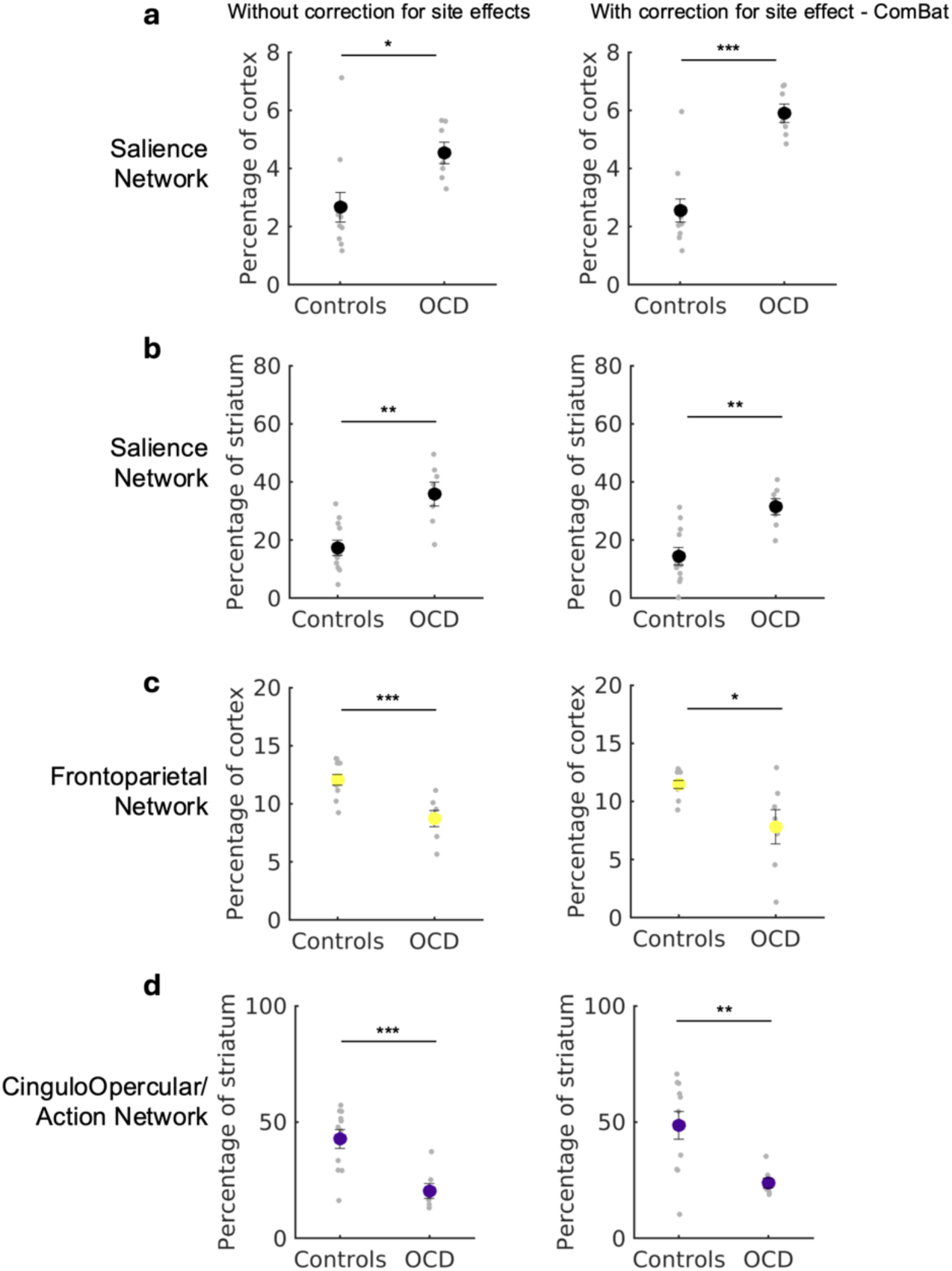
Differences in in networks expansion and contractions replicated when correcting for site effects. The data in the present study is drawn from different sites, which use different imaging sequences and MRI machines. To evaluate if effects of site or scanner impact contribute to the observed patterns of expansion and contractions in individuals with OCD, we applied ComBat harmonization (Fortin et al., Neuroimage 2018) to functional network topography (size) and repeated our analysis. The original (unadjusted) values were used for one of the healthy controls (MyConnectome) because they were the only individuals from their respective sites. Left column: results without ComBat harmonization (reported in the main text); Right column: results with ComBat harmonization to account for potential site effects. Results of salience network expansion in individuals with OCD versus healthy controls remained statistically significant even when accounting for potential site effects both in **a,** cortical (non-parametric Wilcoxon rank-sum t-test, P < 0.001, rank sum =101, Cohen’s d = 2.8) and **b,** striatal (non-parametric Wilcoxon rank-sum test, P = 0.003, rank sum =98, Cohen’s d = 1.87) regions. Similarly, we observed **c**, cortical contraction of the Frontoparietal network (shown in yellow) (non-parametric Wilcoxon rank-sum t-test, P = 0.03, rank sum = 42, Cohen’s d = - 1.43) and **d,** striatal contraction of the Cingulo-Opercular /Action Network (shown in purple) (non-parametric Wilcoxon rank-sum t-test, P = 0.006, rank sum = 37, Cohen’s d = −1.55) regardless of correction for site effects with ComBat harmonization.

**Extended Data Fig. 4.**
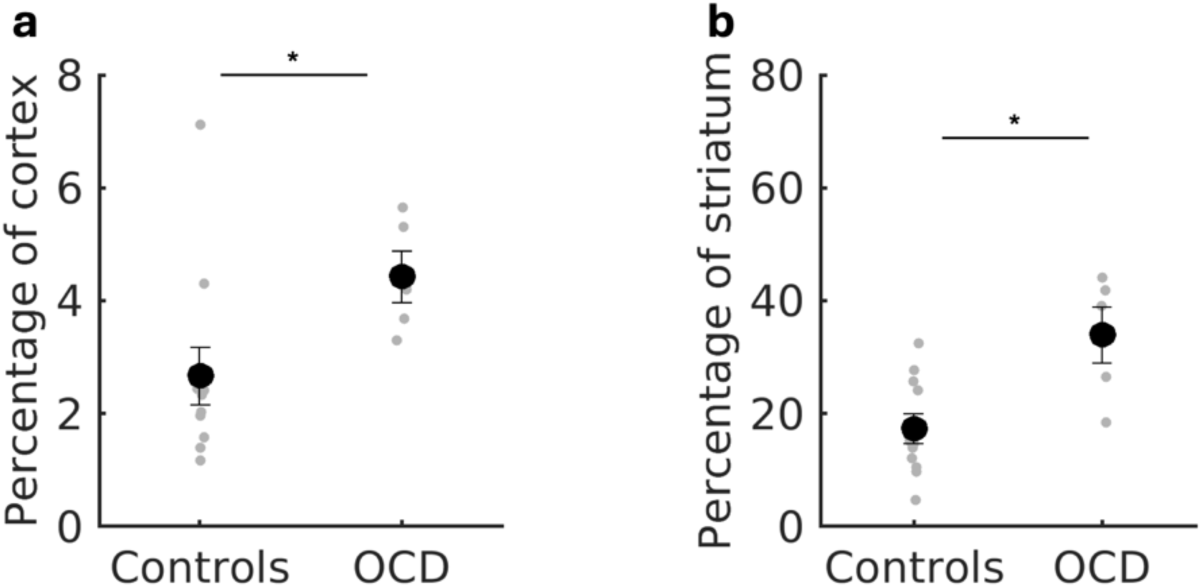
Cortical and striatal salience network expansion in OCD is not driven by individuals with a comorbid diagnosis. **a,** Cortical expansion of the salience network in OCD remained statistically significant even when excluding individuals with a comorbid diagnosis as determined by a non-parametric Wilcoxon rank-sum t-test (P=0.027, rank sum = 62, Cohen’s d = 1.15). In this reduced sample of OCD patients, on average, the salience network occupied 66% more of the cortical surface relative to the mean in healthy controls (4.43% ± 1.01% of cortex in OCD versus 2.67% ± 1.69% of cortex in healthy individuals). This difference gave rise to a large group-level effect. **b,** Similarly, striatal expansion of the salience network in OCD remained statistically significant even when excluding individuals with a comorbid diagnosis as determined by a non-parametric Wilcoxon rank-sum t-test (P=0.013, rank sum = 64, Cohen’s d = 1.78; 34% ± 11.06% of striatum in OCD versus 17.36% ± 8.75% of striatum in healthy individuals). Data are presented as mean ± standard deviation.

**Extended Data Fig. 5.**
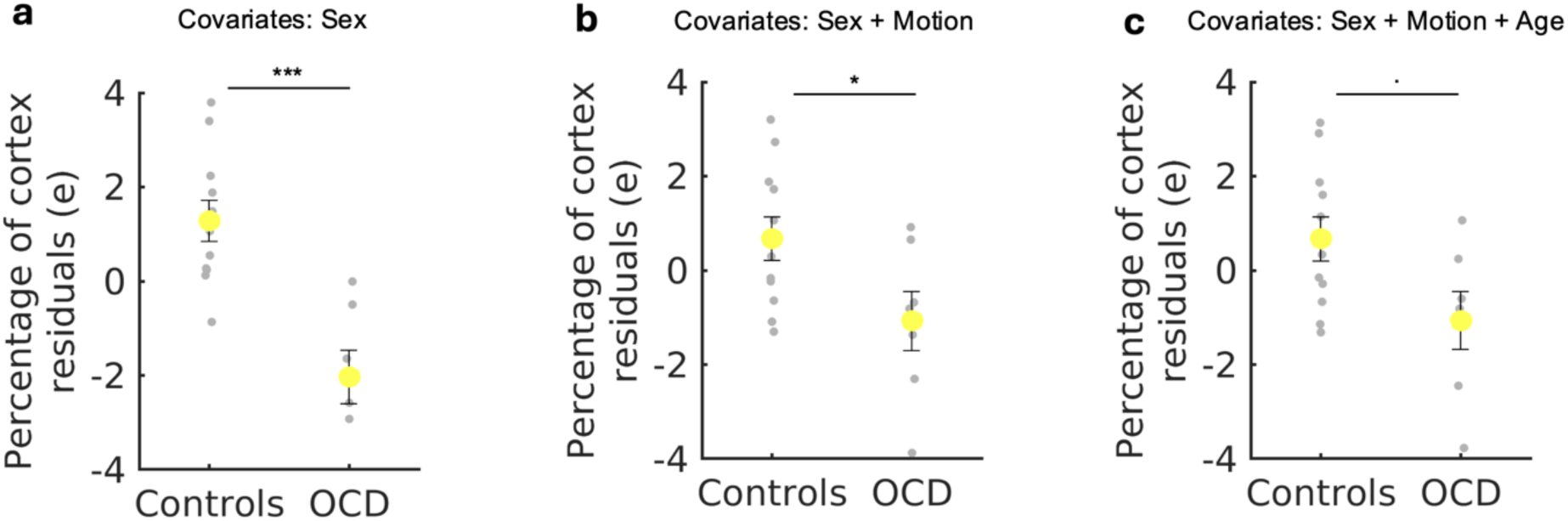
Cortical contraction of the frontoparietal network when controlling for sex ratio, and individual differences in head motion and age. **a,** The size of the frontoparietal network was regressed against sex, and group comparisons were repeated using the residuals (e). Contraction of the frontoparietal network in OCD individuals compared to healthy controls remained statistically significant (non-parametric Wilcoxon rank-sum t-test, P < 0.001, rank sum = 30). **b,** This analysis was repeated with head motion (measured as the percentage of volume retained after motion censoring) as additional covariate and differences between OCD individuals and healthy controls remained significant (non-parametric Wilcoxon rank-sum t-test, P = 0.044, rank sum = 44). **c,** When also age was included as part of the set of covariates, the results narrowly missed reaching statistical significance (non-parametric Wilcoxon rank-sum t-test, P = 0.069, rank sum = 46). Data are presented as mean ± standard deviation.

**Extended Data Fig. 6.**
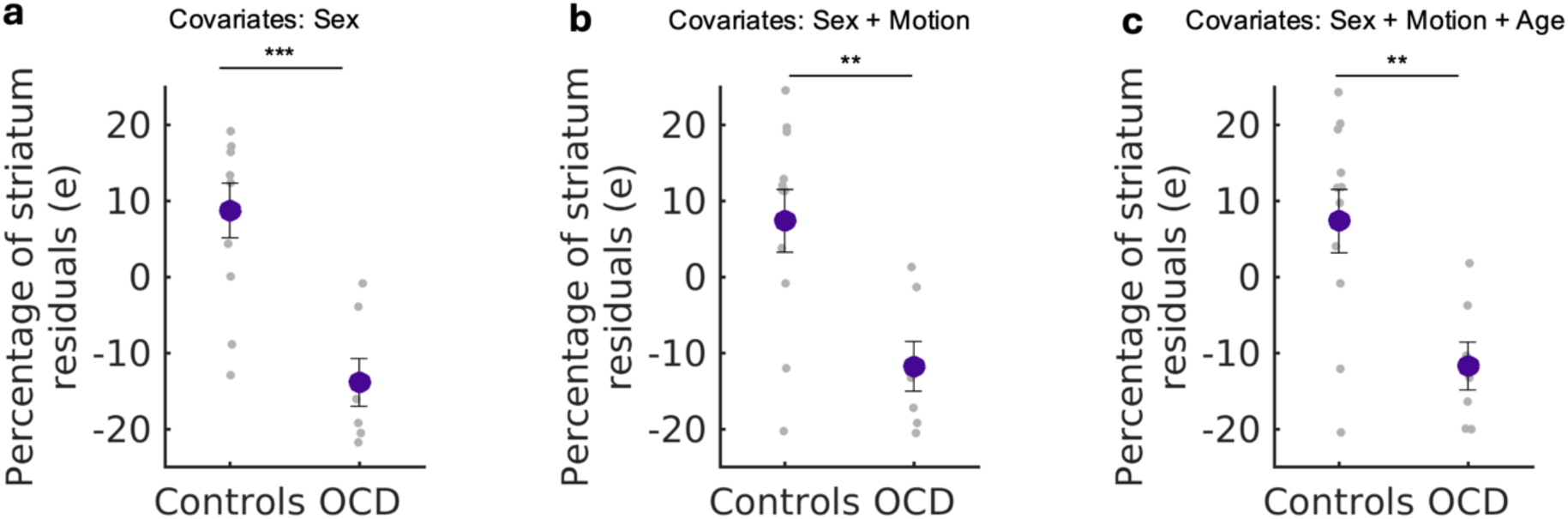
Striatal contraction of the cingulo-opercular network when controlling for sex ratio, and individual differences in head motion and age. The size of the cingulo-opercular network was regressed against sex, and group comparisons were repeated using the residuals (e). Striatal contraction of the cingulo-opercular network in individuals with OCD compared to healthy controls remained statistically significant cortical (non-parametric Wilcoxon rank-sum t-test, P = 0.001, rank sum = 32). **b-c**, This analysis was repeated with head motion (measured as the percentage of volume retained after motion censoring) and age (in years) as additional covariates. In all models, the cingulo-opercular network in the striatum remained significantly smaller in OCD individuals (Sex + Motion: Wilcoxon rank-sum t-test, P = 0.008, rank sum = 38; Sex + Motion + Age: Wilcoxon rank-sum t-test, P = 0.011, rank sum = 39). Data are presented as mean ± standard deviation.

**Extended Data Fig. 7.**
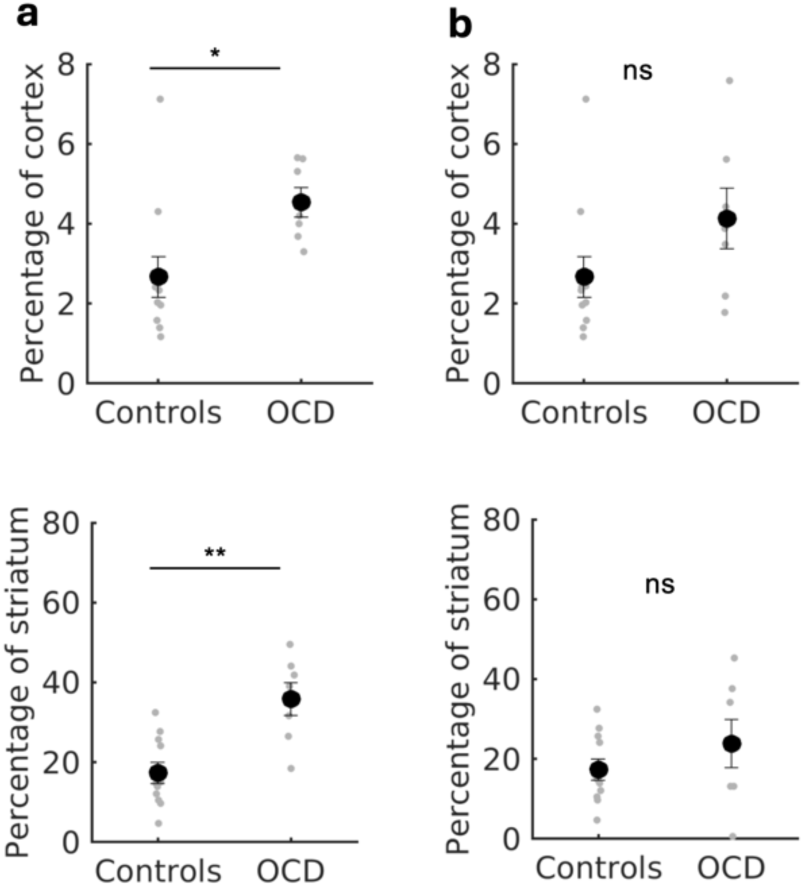
Frontostriatal salience network expansion can’t be detected with low amount of data. **a,** Group differences between OCD and healthy controls in frontostriatal salience network expansion as reported in the main text. **b,** We repeated our pipeline on a reduced dataset including only 1h multi-echo fMRI data per-subject, In this analysis, we could not detect a difference between OCD and control with respect to the cortical (non-parametric Wilcoxon rank-sum t-test, P=0.104, rank sum = 85) and striatal (non-parametric Wilcoxon rank-sum t-test, P=0.425, rank sum =76) size of the salience network. This might explain why previous reports, including only 1h multi-echo fMRI data per-subject, failed to find a difference between healthy controls and OCD patients. We also show that more than 1h of high-quality fMRI data needed per-subject for reliable mapping of salience network, and consistent detection of expansion in individuals.

